# Induction of transmucosal protection by oral vaccination with an attenuated *Chlamydia*

**DOI:** 10.1101/2023.01.30.526385

**Authors:** Yihui Wang, Rongze He, Halah Winner, Marie-Claire Gauduin, Nu Zhang, Cheng He, Guangming Zhong

## Abstract

*Chlamydia muridarum* has been used to study chlamydial pathogenesis since it induces mice to develop hydrosalpinx, a pathology observed in *C. trachomatis*-infected women. We identified a *C. muridarum* mutant that is no longer able to induce hydrosalpinx. In the current study, we evaluated the mutant as an attenuated vaccine. Following an intravaginal immunization with the mutant, mice were protected from hydrosalpinx induced by wild type *C. muridarum*. However, the mutant itself productively colonized the mouse genital tract and produced infectious organisms in vaginal swabs. Nevertheless, the mutant failed to produce infectious shedding in the rectal swabs following an oral inoculation. Importantly, mice orally inoculated with the mutant mounted transmucosal immunity against challenge infection of wild type *C. muridarum* in the genital tract. The protection was detected as early as day 3 following the challenge infection and the immunized mice were protected from any significant pathology in the upper genital tract. However, the same orally immunized mice failed to prevent the colonization of wild type *C. muridarum* in the gastrointestinal tract. The transmucosal immunity induced by the oral mutant was further validated in the airway. The orally vaccinated mice were protected from both lung infection and systemic toxicity caused by intranasally inoculated wild type *C. muridarum* although the same mice still permitted the gastrointestinal colonization by the wild type *C. muridarum*. These observations suggest that the mutant *C. muridarum* may be developed into an <u>intr</u>acellular <u>o</u>ral <u>v</u>accine vector (or IntrOv) for selectively inducing transmucosal immunity in extra-gut tissues.

## Introduction

Sexually transmitted infection with the obligate intracellular bacterium species *Chlamydia trachomatis* can lead to inflammatory pathologies in the upper genital tract, potentially resulting in pelvic inflammatory diseases and infertility (1, 2). An effective vaccine against *C. trachomatis* is considered necessary for preventing the sequelae because *C. trachomatis* infection is asymptomatic or lacks specific symptoms. However, despite extensive research and development efforts, there is still no licensed Chlamydia vaccine for humans (3). When chlamydial organisms were successfully isolated from human samples more than 70 years ago (4, 5), intramuscular injection of formalin-inactivated chlamydial organisms was tested for preventing trachoma caused by *C. trachomatis* in school children (6–8). Unfortunately, the inactivated whole organism vaccines not only failed to induce lasting protection but also exacerbated conjunctivitis in some cases. The failed trachoma vaccine trials motivated chlamydial researchers to search for subunit vaccines. Over the years, many chlamydial components have been evaluated as vaccine candidates in various preclinical models (9, 10). A recent study has demonstrated that the inactivated chlamydial organisms may be modified to induce protective immunity with no detrimental effects in preclinical models (11). However, none of the above has ever been advanced to efficacy evaluations in humans although a phase I safety evaluation has been carried out for one of the candidates (12).

Mouse infection with *Chlamydia muridarum* is one of the most extensively used preclinical models for investigating chlamydial pathogenesis and evaluating chlamydial vaccines (9, 13–17). *C. muridarum* is known to cause hydrosalpinx and infertility in mice following intravaginal inoculation (18–20), which mimics the tubal adhesion/infertility observed in women under laparoscopy (2, 21, 22). The fact that genital *C. muridarum* infection is self-limited and often ends up with oviduct scaring suggests that there may be a partial fitness between *C. muridarum* and the female mouse genital tract mucosal tissue. In contrast, intranasal inoculation with *C. muridarum* leads to severe acute pneumonia or death (23, 24), suggesting that there is a lack of fitness between *C. muridarum* and the mouse airway mucosal tissue. Thus, *C. muridarum* can be used for evaluating immunity against chlamydial infection under different infection conditions (25, 26).

Besides infecting the mucosal tissues in the mouse genital tract and airway, *C. muridarum* is also known to colonize the gastrointestinal (GI) tract for a long period of time (27, 28). Interestingly, various chlamydial species have been detected in their corresponding host GI tracts (29–33). Although the significance of chlamydial species in the GI tract remains unknown, recent studies based on the models of *C. muridarum* interaction with mouse mucosal tissues have revealed that the order of chlamydial exposure to mouse tissue sites may significantly influence the consequence of the gut *C. muridarum* (34, 35). When a naïve mouse is exposed to *C. muridarum* in the GI tract first, *C. muridarum* may function as an oral vaccine. It has been reported that oral inoculation with *C. muridarum* can induce robust transmucosal immunity against subsequent chlamydial infections in both the genital tract (25) and airway (26), suggesting that *C. muridarum* may be developed into an oral vaccine for inducing transmucosal immunity. To further improve the safety of oral Chlamydia vaccines, efforts have been made to evaluate various genital tract pathogenicity-attenuated *C. muridarum* mutants (36, 37).

We recently identified a mutant *C. muridarum* that is highly attenuated in inducing pathogenicity in the female mouse genital tract following an intravaginal inoculation (38, 39). In the current study, we compared the genital pathogenicity-attenuated mutant for its ability to induce protection in the genital tract following intravaginal immunization versus oral immunization. Intravaginal immunization with the mutant protected mice from developing hydrosalpinx induced by wild type *C. muridarum*. However, the protection was accompanied with productive colonization of the mutant itself in the mouse genital tract, which led to the release of infectious organisms in vaginal swabs. In contrast, oral inoculation with the mutant failed to produce infectious shedding in the rectal swabs. More importantly, orally immunized mice developed robust transmucosal immunity against challenge infection by wild type *C. muridarum* in the genital tract although these mice permitted the colonization of wild type *C. muridarum* in the GI tract. The oral mutant-induced selective protection in the extra-gut tissues but not the gastrointestinal tract was further validated in a mouse airway model in which mice were intranasally challenged with wild type *C. muridarum*. Thus, the mutant *C. muridarum* may be developed into an <u>intr</u>acellular <u>o</u>ral <u>v</u>accine vector (or IntrOv) for selectively inducing transmucosal immunity.

## Results

### 1. A genital pathogenicity-attenuated *C. muridarum* mutant induces protection against subsequent infection by wild type *C. muridarum*

Using a Pasteur passage selection scheme (38), we previously identified a *C. muridarum* mutant (CMmut, clone CMG28.51.1) that was no longer able to induce hydrosalpinx following an intravaginal inoculation (39). Intravaginal inoculation with CMmut was evaluated for inducing protective immunity against subsequent challenge infection by wild type *C. muridarum* (CMwt). As shown in Fig. 1, in the sucrose-phosphate-glutamate buffer (SPG)-treated mice (mock immunization control), intravaginal challenge infection with CMwt produced a typical one-month live chlamydial organism shedding course from the genital tract. All mice cleared infection by day 35. However, the CMwt challenge infection-induced shedding course was significantly shortened in the CMmut-immunized mice. All mice cleared infection by day 21. The shedding levels in terms of infectious forming units (IFUs) per swab were significantly reduced on days 3, 7 & 14. Clearly, prior genital tract exposure to CMmut was sufficient for inducing anti-chlamydial immunity in the female genital tract. Further, the CMmut-induced protective immunity significantly protected mice from developing hydrosalpinx induced by CMwt. Only one of the 8 mice in the immunized group developed hydrosalpinx while 6 of 8 control mice did so. The immunized group only had a mean hydrosalpinx score of 0.25 while the control group had a score of 4.5. These results have demonstrated that although intravaginal inoculation with the CMmut clone itself can no longer induce hydrosalpinx (39), CMmut can still induce protective immunity, suggesting that it can be developed into an attenuated vaccine. However, the CMmut-induced protection was accompanied with its own colonization, resulting in the release of live CMmut organisms from the genital tract, which raises a safety concern on the potential spreading of the vaccine strain in the vaccinated host population.

**Fig.1.**
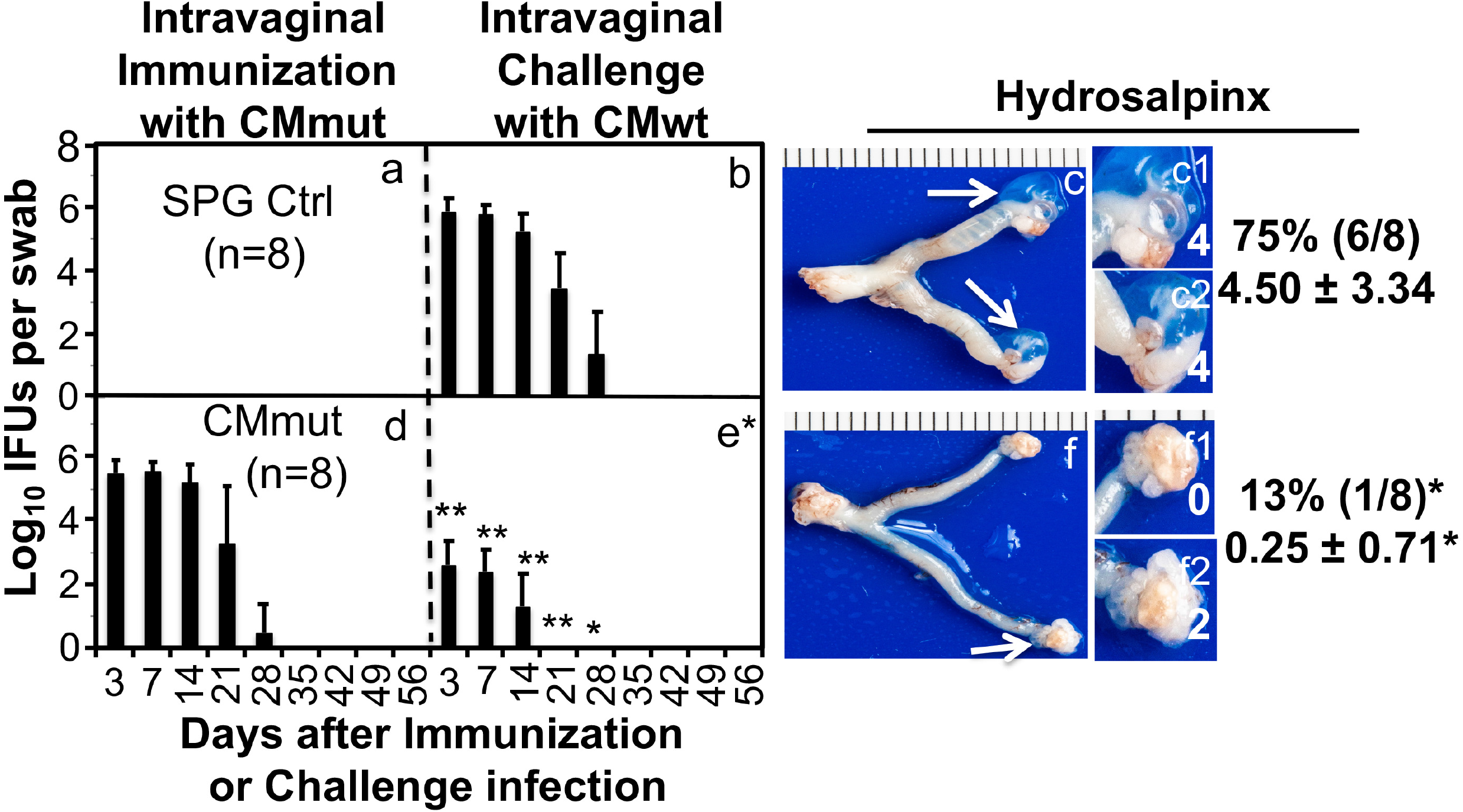
Effect of intravaginal immunization with an attenuated *Chlamydia* on subsequent challenge infection by wild type *Chlamydia*. Groups of female C57BL/6J mice were intravaginally inoculated with SPG buffer (Ctrl, n=8, panel a) or a mutant *C. muridarum* (CMmut at an inoculum dose of 2 x 10^5^ inclusion forming units or IFUs, n=8, d) as immunization for 56 days. Subsequently, both groups of mice were intravaginally challenged with wild type *C. muridarum* (CMwt at an inoculum dose of 2 x 10^5^ IFUs, b & e). All mice were monitored for live *Chlamydia* burdens in the genital tract by taking vaginal swabs on days 3, 7 and weekly thereafter (X-axis) following the immunization (panels a & d) and challenge infection (b & e) respectively. The number of live organisms recovered from each vaginal swab was expressed as log_10_IFUs (Y-axis). 56 days after the challenge infection, all mice were sacrificed for observing hydrosalpinx (c & f). Only one representative image of the entire genital tract was shown for each group. Oviducts positive for hydrosalpinx were marked with white arrows. The magnified images of oviduct/ovary regions (with hydrosalpinx scores indicated in white numbers) were shown on the right of the overall genital tract image. Both the hydrosalpinx incidence (along with group sample size) and severity score from each group were listed next to the corresponding group images. *p<0.05, **p<0.01 (Fisher’s Exact for comparing incidences while Wilcoxon for scores or IFUs). Data were from 2 or 3 independent experiments. Note that intravaginal immunization with CMmut induced significant protection against both infection and pathogenicity of CMwt.

### 2. Mice orally inoculated with CMmut fail to release live chlamydial organisms

We previously reported that CMmut failed to spread from the genital tract into the gastrointestinal (GI) tract (40) although CMwt readily did so (41), suggesting that CMmut might not produce infectious shedding in the GI tract. As shown in Fig. 2, following a direct oral inoculation with 1 x 10^7^ IFUs of CMmut, mice failed to release any live chlamydial organisms from either the rectal or vaginal swabs while mice orally inoculated with CMwt continuously shed live chlamydial organisms from the GI tract. Further, following an intracolon inoculation, only a minimal level of CMmut live chlamydial organisms was detected in the rectal swabs and all CMmut-inoculated mice clear infection by day 14. However, the intracolon-inoculated CMwt maintained a steady level of live organism shedding. These observations indicate that CMmut can be rapidly cleared from the mouse colon, which may partially explain why oral CMmut failed to shed live chlamydial organisms in the rectal swabs. Thus, CMmut may not cause spreading when it is delivered orally.

**Fig.2.**
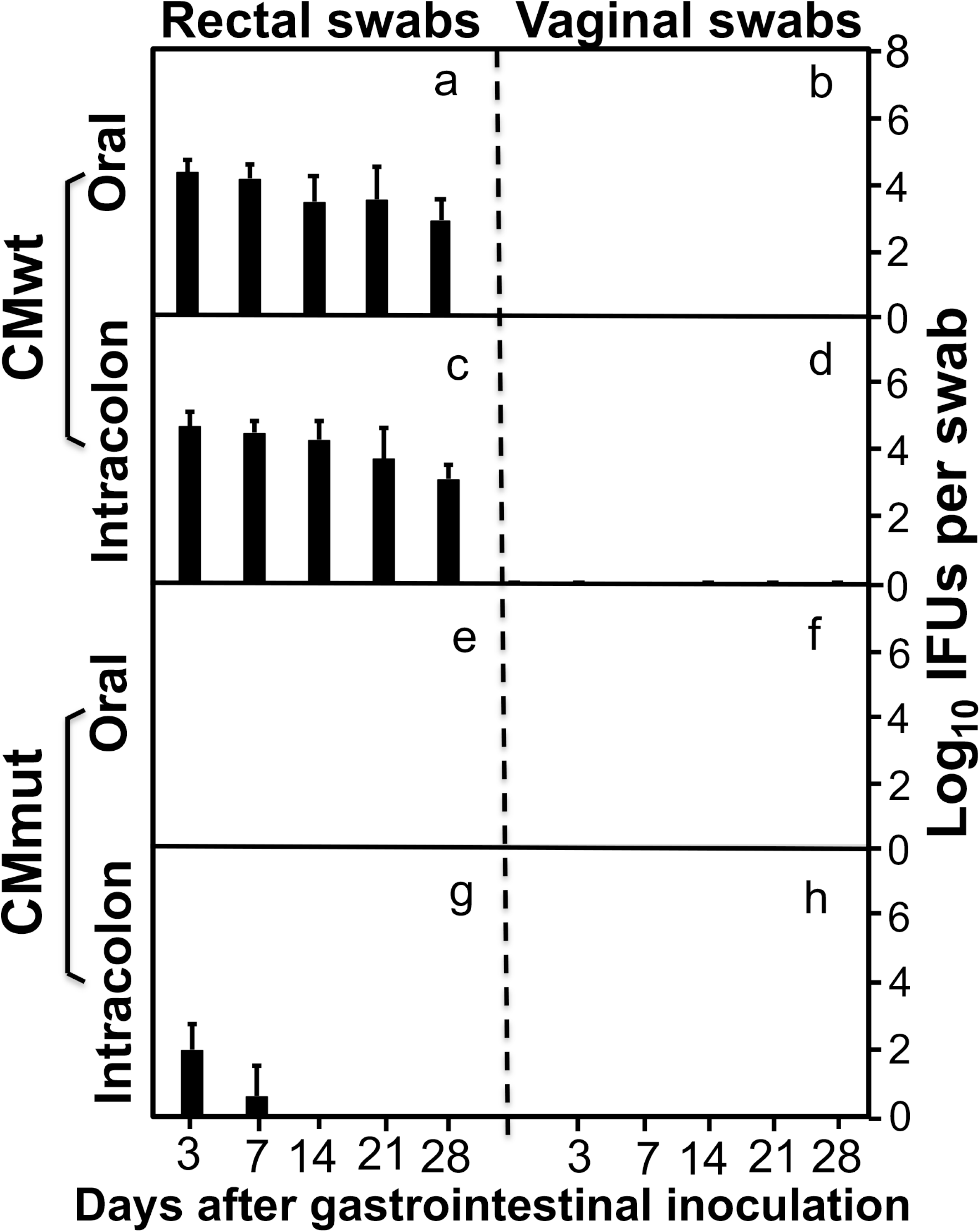
Comparison of live chlamydial organism shedding courses between mice orally or intrarectally inoculated with wild or mutant *Chlamydia*. Groups of female C57BL/6J mice (n=5) were either orally (panels a, b, e & f) or intracolonally (c, d, g & h) inoculated with wild type *Chlamydia* (CMwt at an inoculum dose of 2 x 10^5^ IFUs, a-d) or mutant *Chlamydia* (CMmut at an inoculum dose of 1 x 10^7^ IFUs, e-h). All mice were monitored for live *Chlamydia* burdens in the gastrointestinal tract and genital tract by taking rectal and vaginal swabs on days 3, 7 and weekly thereafter (X-axis). The number of live organisms recovered from each swab was expressed as log_10_IFUs (Y-axis). Data were from 2 experiments. Note that orally inoculated CMmut failed to shed live chlamydial organisms from either gastrointestinal tract or genital tract although minimal shedding was detected following intrarectal inoculation.

### 3. Oral immunization with CMmut protects against infection and pathogenicity of CMwt in the genital tract

Since oral CMmut failed to shed live organisms, oral CMmut was further evaluated for the induction of protective immunity in the genital tract. As shown in Fig. 3, mice orally immunized with CMmut significantly reduced the shedding levels of CMwt live organisms in the genital tract. The SPG mock-immunized control group developed a typical one month long shedding course caused by intravaginal challenge infection with CMwt while the CMmut-immunized group significantly shortened the shedding course of CMwt. Live CMwt organisms were only detected on days 3 & 7 with significantly reduced levels at both time points in the oral CMmut-immunized mice. These observations have demonstrated that oral CMmut is able to induce transmucosal immunity in the female genital tract. It is worth noting that a steady level of live organism shedding was detected in the rectal swabs of the CMmut-immunized mice following intravaginal challenge infection with CMwt. The rectal live organisms might be caused by the intravaginally challenged CMwt since CMwt is known to spread from the genital tract into the GI tract (41). More importantly, the oral CMmut-induced transmucosal immunity was sufficient for preventing pathology in the upper genital tract (Fig. 4). None of the oral CMmut-immunized mice developed hydrosalpinx following intravaginal challenge infection with CMwt. The lack of hydrosalpinx in the immunized mice was validated by examining oviduct dilation under microscopy. Only two of the 6 immunized mice developed oviduct dilation with a mean dilation score of 0.33 while all control mice developed significant oviduct dilation with a mean score of 4.57. The above observations have demonstrated that oral immunization with CMmut is an efficient approach for inducing transmucosal immunity in the female genital tract.

**Fig.3.**
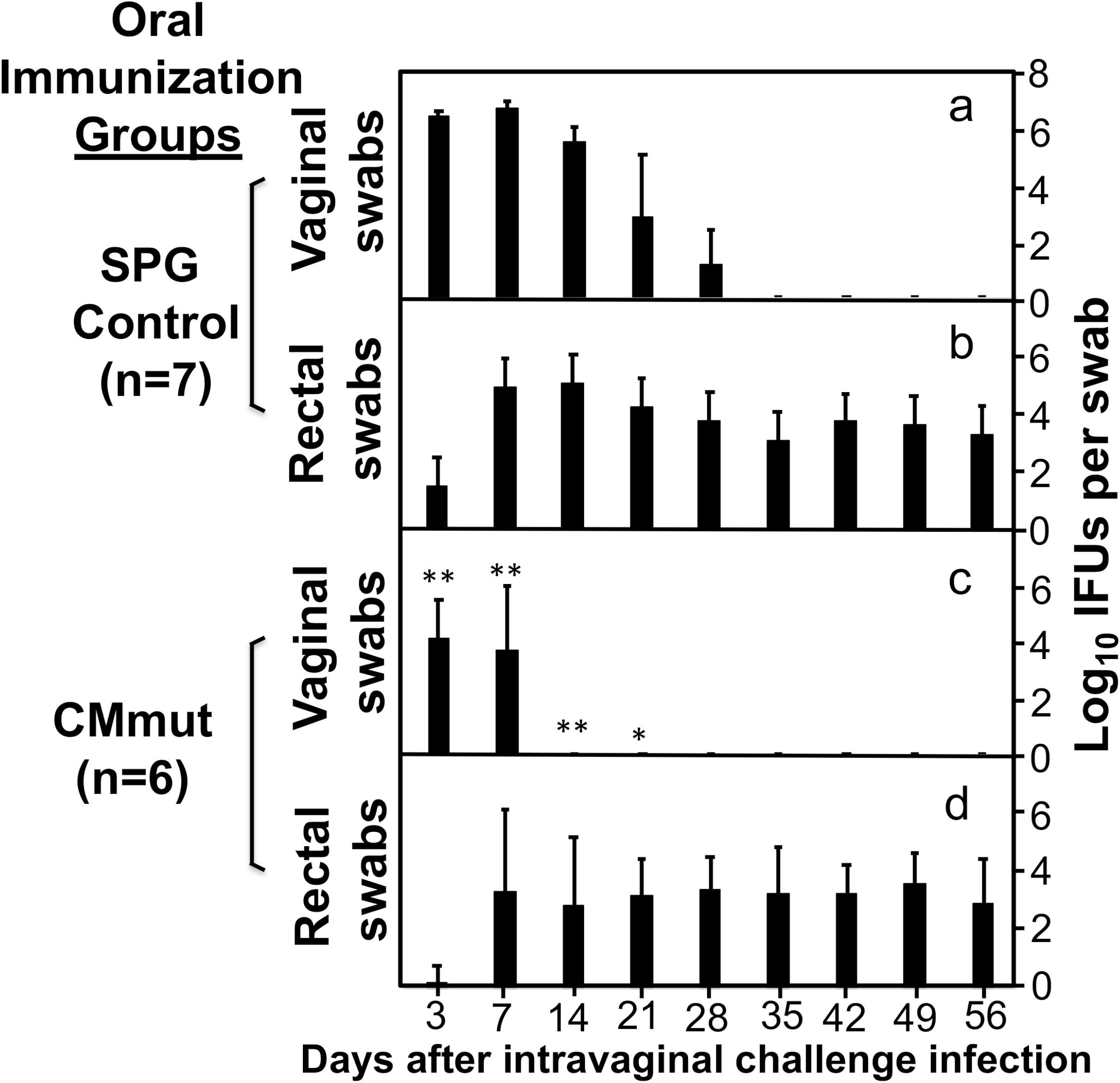
Effect of oral immunization with attenuated *Chlamydia* on subsequent challenge infection by wild type *Chlamydia* in the genital tract. Female C57BL/6J mice orally inoculated with SPG (control, panels a & b, n=7) or mutant *C. muridarum* (CMmut at an inoculum dose of 1 x 10^7^ IFUs, c & d, n=6) as immunization for 28 days. Both groups of mice were then intravaginally challenged with 2 x 10^5^ IFUs of wild type *C. muridarum* (CMwt). All mice were monitored for live chlamydial organisms from both the genital tract (vaginal swabs, a & c) and gastrointestinal tract (rectal swabs, b & d) on days 3, 7 and weekly thereafter following the challenge infection. The results were expressed as Log_10_ IFUs per swab (Y-axis). Data were from 2 different experiments. Note that mice orally immunized with CMmut were protected from genital tract infection by CMwt (*p<0.05, **p<0.01, Wilcoxon for comparing IFUs at different time points between panels a and c).

**Fig.4.**
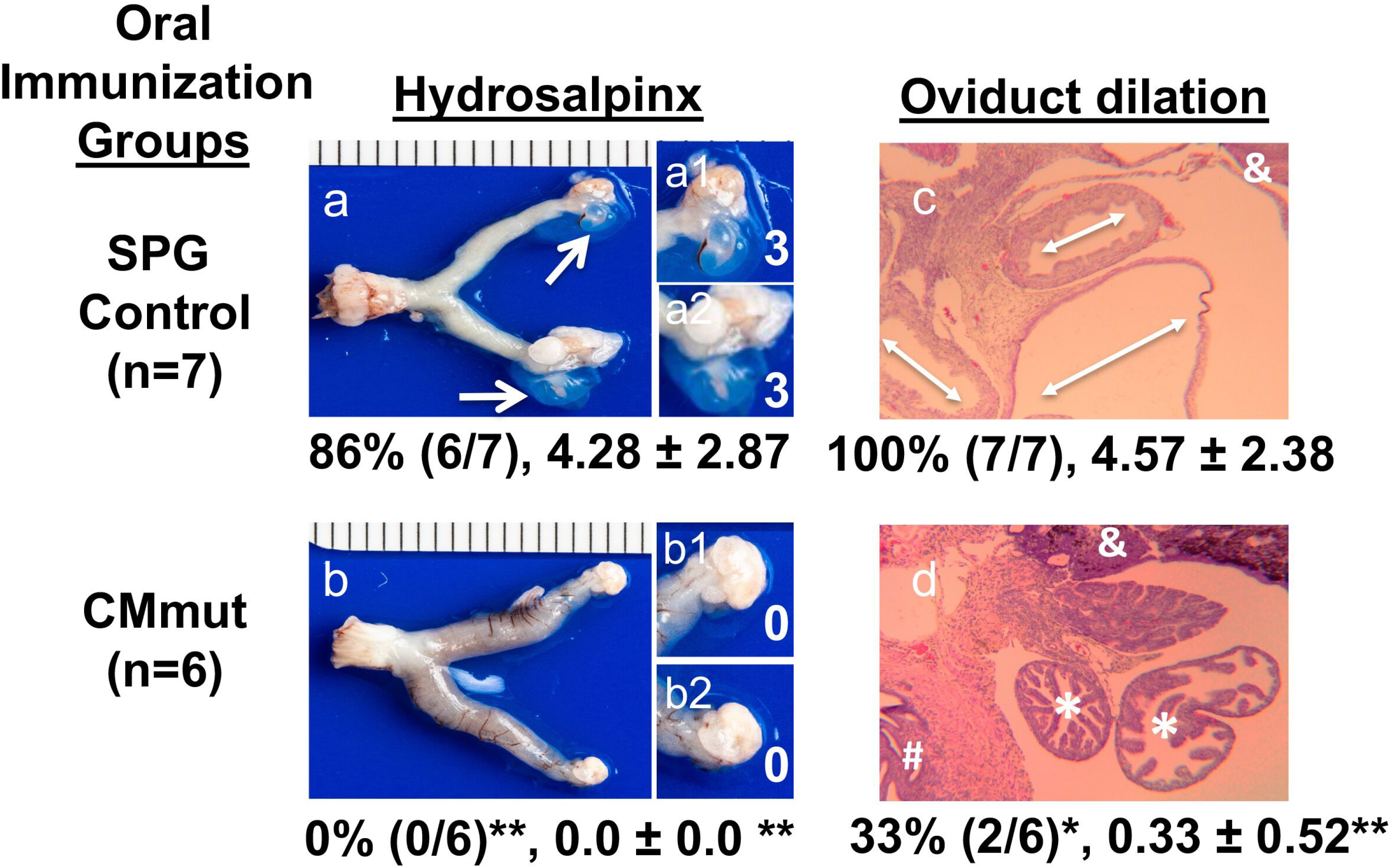
Effect of oral immunization with attenuated *Chlamydia* on pathogenicity of wild type *Chlamydia* in the genital tract. The same groups of C57BL/6J mice orally immunized and intravaginally challenged as described in figure 3 legend were sacrificed on day 56 after the intravaginal challenge infection for observing genital tract pathology. Hydrosalpinx was visually evaluated. Only one representative image of the entire genital tract was shown for each group. Oviducts positive for hydrosalpinx were marked with white arrows. The magnified images of oviduct/ovary regions (with hydrosalpinx scores indicated in white numbers) were shown on the right of the overall image. Both the hydrosalpinx incidence (along with group sample size) and severity score from each group were listed under the corresponding group images. The same excised genital tract tissues were further processed for monitoring oviduct dilation under microscopy. After H&E staining, tissue sections of the genital tissues were first examined for the overall appearance of the oviduct tissues under a 4x objective lens (c & d). Representative normal oviduct cross-section was labeled with a white star “*”, ovary with “&” and uterine horn tissue with “#” while dilated oviducts were indicated with the white double arrowhead arrows. Both hydrosalpinx (a & b) and oviduct dilation (c & d) were semi-quantitatively measured as described the materials and method section. Data were from 2 independent experiments. *p<0.05 & **p<0.01, Fisher’s Exact for comparing incidence rates while Wilcoxon for comparing scores between the control and CMmut-immunized group. Note that CMmut-immunized group showed significantly reduced hydrosalpinx and oviduct dilation.

### 4. Oral CMmut induces transmucosal protection against infection and pathogenicity of CMwt in the airway

Having demonstrated the robust transmucosal protection in the female genital tract, we next tested whether the oral CMmut-induced transmucosal immunity can also be detected in the airway. Intranasal inoculation with CMwt is known to cause mouse pneumonia or death (24), which was used for evaluating transmucosal immunity induced by oral CMmut in the current study. As shown in Fig. 5, in the control group, intranasal challenge infection with CMwt significantly decreased mouse body weight, which is consistent with previous studies (23, 24, 26, 42). However, the oral CMmut-immunized mice gained body weight in the same period following the intranasal challenge infection with CMwt. Thus, oral CMmut is sufficient for protecting mice from the systemic toxicity caused by airway CMwt. When the live CMwt burden was monitored in the mouse lung tissue on day 9 after intranasal inoculation, the CMmut-immunized mice were completely protected from any live chlamydial organisms while the SPG control mice harbored >10 million IFUs of live chlamydial organisms per lung. The difference in CMwt burden in the lung may partially explain the lack of toxicity in the oral CMmut-immunized mice. It is worth noting again that despite the completion protection of the lung, low levels of live chlamydial organisms were detected in the stomach and colon. These live chlamydial organisms may come from the airway CMwt since CMwt is known to establish lasting colonization in the gut following an inoculation elsewhere (43, 44).

**Fig. 5.**
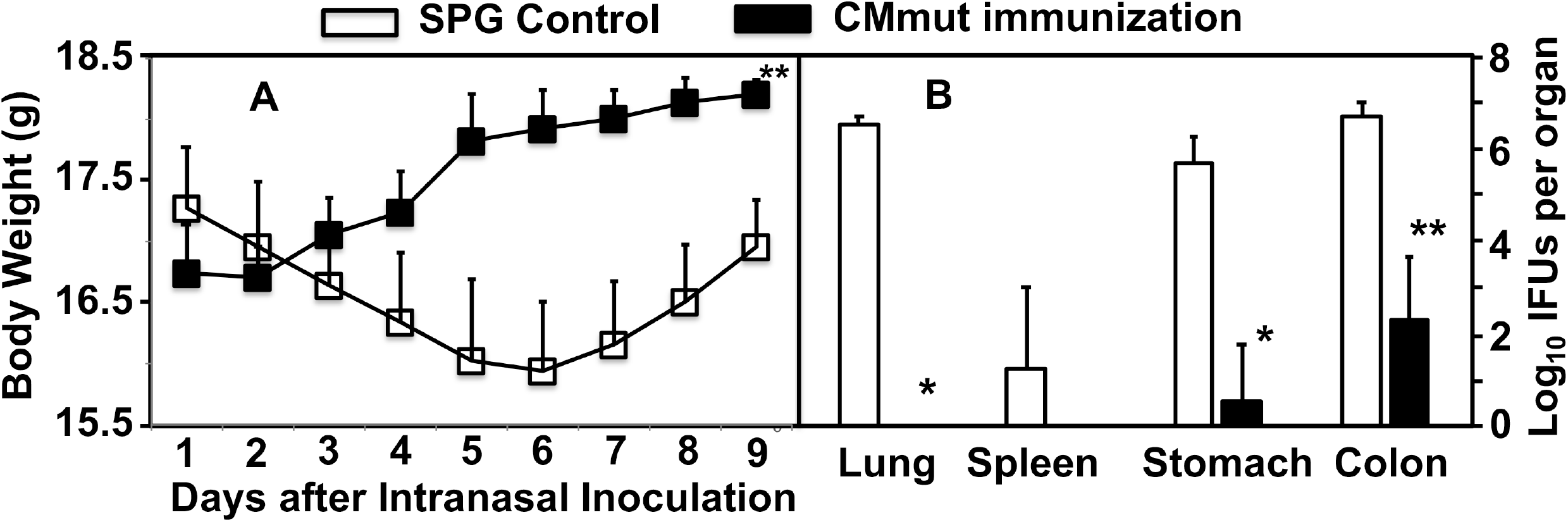
Effect of oral immunization with attenuated *Chlamydia* on subsequent infection by wild type *Chlamydia* in the airway. Female C57BL/6J mice orally inoculated with SPG (control, open square/bar, n=5) or mutant *C. muridarum* (CMmut at an inoculum dose of 1 x 10^7^ IFUs, solid square/bar, n=5) as immunization for 28 days. Both groups of mice were then intranasally challenged with 5,000 IFUs of wild type *C. muridarum* (CMwt). All mice were monitored for body weight daily (A) and the mouse body weight were measured in grams (Y-axis). All mice were sacrificed for measuring live chlamydial organism burdens in lungs, spleen, stomach and colon tissues on day 9 after intranasal infection (B). The results were expressed as Log_10_ IFUs per organ (Y-axis). Data were from 2 different experiments. *p<0.05 & **p<0.01, Wilcoxon for comparing body weight (area-under-curve) or IFUs between SPG group and CMmut immunization group. Note that mice orally immunized with CMmut were protected from airway infection by CMwt.

## Discussion

There is an urgent need to develop an effective vaccine against *C. trachomatis* in humans. It is obvious that an ideal Chlamydia vaccine would be a subunit vaccine. However, years of extensive efforts in evaluating hundreds of chlamydial components as vaccine candidates have yielded no licensable vaccines. Perhaps, it is time to explore or re-explore non-subunit vaccine approaches. A recent study has revealed that inactivated whole chlamydial organisms can be modified to induce protective immunity without exacerbating pathology (11), which is similar to the immunity induced by live chlamydial infection. Using the *C. muridarum* model, it has been shown that inoculation with live *C. muridarum* induces the strongest protection against subsequent challenge infection (45–47). However, inoculating live chlamydial organisms into either the genital tract or the airway causes pathologies. Instead, we have recently shown that oral inoculation with *C. muridarum* is not only non-pathological but also induces strong transmucosal protection in extra-gut tissues (25, 26). These observations suggest that it is possible to develop a safe and effective oral Chlamydia vaccine. To further improve the safety of oral Chlamydia vaccines, various genital tract pathogenicity-attenuated *C. muridarum* mutants have been evaluated as oral vaccines (36, 37).

In the current study, we compared the genital tract pathogenicity-attenuated *C. muridarum* mutant or CMmut (clone G28.51.1) for its ability to induce protection against infection with wild type *C. muridarum* (CMwt) following either an intravaginal immunization or oral immunization. The results have led us to conclude that oral immunization is a more favorable approach. First, although either intravaginal or oral immunization route was sufficient for CMmut to induce protection against genital infection and pathogenicity caused by CMwt, intravaginal CMmut itself produced significant shedding of infectious organisms from the immunized mice while oral CMmut failed to do so. Thus, oral immunization may minimize the spreading of the vaccine strain in the vaccinee population. Second, intravaginal inoculation is obviously more difficult to administer and may be hard to be accepted. Thus, the compliance rate will be a major concern. On the contrary, oral vaccination is easy to administer and has been successfully used for other vaccines. These benefits have motivated us to propose to develop the CMmut clone G28.51.1 into an intracellular oral vaccine or IntrOv for short. Finally, the genital tract protection induced by oral CMmut is based on transmucosal immunity. The robust transmucosal immunity was also validated in the airway. Thus, the clone G28.51.1 may also be developed into an intracellular oral vector for delivering prevention or intervention reagents against non-chlamydial diseases in extra-gut tissues.

Although oral CMmut- or IntrOv-induced immunity efficiently protected both the mouse genital and airway mucosal tissues from infection and pathogenicity caused by CMwt, the immunity was insufficient for protecting the same mouse’s GI tract against colonization by CMwt. First, the IFUs detected in the GI tract were not caused by the IntrOv organisms since oral inoculation with IntrOv did not produce any IFUs in either the rectal or vaginal swabs. Second, the GI IFUs detected in mice challenged with CMwt might be due to the spreading of genital or airway CMwt organisms into the GI tract. This is because it has been demonstrated that any mucosal inoculation with *C. muridarum* can always lead to systemic spreading (43) and the systemic *C. muridarum* can only establish long-lasting colonization in the GI tract (44). By following the spreading of genital *C. muridarum* into the GI tract (41), two distinct and complementary pathways have been identified for systemic *C. muridarum* to home to the GI tract (48). The primary pathway is the spleen-stomach pathway while the secondary pathway is the liver-intestinal route. Nevertheless, the precise molecular and cellular basis of *C. muridarum* trafficking into the GI tract remains unclear although dendritic cells (DCs) have been proposed to play an essential role (49).

The next question is why the oral IntrOv-induced immunity is effective against chlamydial infections in the mucosal tissues of both the genital tract and airway but not the GI tract.

Using a combination of knockout mice and mutant *C. muridarum* (the IntrOv clone), we have recently shown that the long-lasting colonization of *C. muridarum* in mouse colon is dependent on its ability to evade colonic IFNγ that is produced by the group 3 innate lymphoid cells (ILC3s) or ex-ILC3s but not other ILCs or conventional lymphocytes (50, 51). Although IFNγ can be produced by many different types of cells, it is likely that *C. muridarum-infected* cells in the GI tract may only be accessible by ex-ILC3s but not other lymphocytes while the *C. muridarum-infected* cells in extra-gut tissues may be accessed by different lymphocytes (34). Oral *C. muridarum* is known to induce transmucosal immunity that is dependent on conventional lymphocytes (25). Thus, we hypothesize that oral IntrOv may also induce conventional lymphocyte-dependent immunity that can only transmucosally prevent *C. muridarum* infection in the female genital tract and airway but not the GI tract. Testing of this hypothesis is underway.

The final but important question is whether the *C. muridarum* model-based findings described in the current study can be used to develop a *C. trachomatis* vaccine for humans. A direct approach is to make the same attenuation mutations in the genome of *C. trachomatis* and develop the mutant *C. trachomatis* clone into an oral vaccine. The two key mutations of IntrOv are loss of function mutations in *tc0237* & *tc0668* respectively (38, 39). The corresponding *C. trachomatis* homologs are *tc849* and *tc389* respectively. Genetic approaches for interrupting a gene in the *C. trachomatis* genome are now available (52). Since the mutation is only restricted to two genes unlike other attenuated *C. muridarum* clones with multiple mutations (36), making the corresponding *C. trachomatis* mutant based on IntrOv described in the current study should not be a problem. Although plasmid-free or plasmid gene deletion *C. muridarum* mutants have also been evaluated as oral vaccine candidates (37), plasmid gene-based mutants may be less stable than chromosomal gene mutants. Production of a stable *C. trachomatis* mutant clone with loss of function mutations in *ct849* and *tc398* may allow us to test the hypothesis that oral delivery of an attenuated *C. trachomatis* mutant may be sufficient for inducing transmucosal immunity in the genital tract.

## Materials and Methods

### 1. Chlamydial organism growth

All *Chlamydia muridarum* (CM) clones used in the current study were derived from strain Nigg3 (Genbank accession# CP009760.1), including a wild type clone CMG13.32.1 (CMwt) and a genital pathogenicity-attenuated clone CMG28.51.1 (CMmut or IntrOv; ref; (38, 39). All chlamydial organisms were propagated in HeLa cells and purified as elementary bodies (EBs) as reported previously (41, 53). Aliquots of the purified EBs were stored at −80°C until use.

### 2. Mouse immunization and challenge infection

Mouse experiments were carried out in accordance with the recommendations in the Guide for the Care and Use of Laboratory Animals of the National Institutes of Health. The protocol was approved by the Committee on the Ethics of Laboratory Animal Experiments of the University of Texas Health Science Center at San Antonio.

Purified CMmut/intrOv or CMwt EBs were used to inoculate six to seven week-old female C57BL/6J mice (000664, Jackson Laboratories, Inc., Bar Harbor, Maine) intragastrically (for CMmut/IntrOv as oral immunization route), intravaginally (as both immunization and challenge infection), intracolonally or intranasally as described previously (20, 24, 26, 41, 54, 55). Five days before intravaginal inoculation, each mouse was injected with 2.5 mg subcutaneous medroxyprogesterone (Depo-Provera; Pharmacia Upjohn, Kalamazoo, MI) suspended in sterile phosphate-buffered saline (PBS). The inoculation dose for oral or intracolon intrOv was 1 x 10^7^ inclusion forming units (IFUs; ref: (50, 56)) while for intranasal CMwt was 5,000 IFUs (based on pre-titration in a pilot experiment). The dose of the remaining inoculation was kept at 2 x 10^5^ IFUs regardless of the routes and *C. muridarum* clones. Following each inoculation, vaginal and/or rectal swabs as well as organ/tissues were taken at the designated time points for monitoring viable *C. muridarum* colonization as described previously (41, 44, 54).

### 3. Titrating live chlamydial organisms recovered from swabs and tissues

To quantitate live chlamydial organisms in vaginal or rectal swabs, each swab was soaked in 0.5 ml of SPG, vortexed with glass beads, and the chlamydial organisms released into the supernatants were titrated on HeLa cell monolayers in duplicate. For tissue samples, each organ was transferred to a tube containing 2 ml SPG for homogenization using an automatic homogenizer (Omni Tissue Homogenizer, TH115, Kennesaw, GA). The homogenates were briefly sonicated and spin at 3000 rpm for 5 min to pellet remaining large debris. The supernatants were titrated for live *C. muridarum* organisms on HeLa cells. The infected cultures were processed for immunofluorescence assay as described previously (39, 57) and below. Inclusions were counted in five random fields per well under a fluorescence microscope. For wells with less than one IFU per field, entire wells were counted. Wells showing obvious cytotoxicity of HeLa cells were excluded. The total number of IFUs per swab was calculated based on the mean IFUs per view, the ratio of the view area to that of the well, dilution factor, and inoculation volumes. Where possible, a mean IFU/swab was derived from the serially diluted and duplicate samples for any given swab. The total number of IFUs/swab was converted into log_10_, which was used to calculate the mean and standard deviation across mice of the same group at each time point.

### 4. Immunofluorescence assay

For immunofluorescence labeling of *C. muridarum* in HeLa cells, a rabbit antibody (designated as R1604, raised with purified *C. muridarum* EBs) was used as a primary antibody to label *C. muridarum*, which was visualized with a goat anti-rabbit IgG conjugated with FITC (green, cat#111-225-144, Jackson ImmunoResearch Laboratories, INC., West Grove PA). The DNA dye Hoechst 3328 (blue, Sigma-Aldrich, St. Louis, MO) was used to visualize nuclei. The dually labeled samples were used for counting for *C. muridarum* under a fluorescence microscope (IX80, Olympus) equipped with a CCD camera (Hamamatsu).

### 5. Evaluating genital tract gross pathology hydrosalpinx microscopically

On day 56 after intravaginal infection, mice were euthanized for evaluating hydrosalpinx in the upper genital tract. Before the tissues were removed fro mice, an *in situ* gross examination was performed for evidence of hydrosalpinx or any other related abnormalities. The severity of hydrosalpinx was scored based on the following criteria: no hydrosalpinx (0), hydrosalpinx detectable only under microscopic examination (1), hydrosalpinx clearly visible with naked eyes but the size was smaller than the ovary on the same side (2), equal to the ovary on the same side (3) or larger than the ovary on the same side (4). Bilateral hydrosalpinx severity was calculated for each mouse as the summed scores of the left and right oviducts. Hydrosalpinx incidence was calculated as the number of mice with a score of 1 or higher divided by the total number of mice in the group.

### 6. Evaluating oviduct dilation microscopically

After microscopic evaluation of hydrosalpinx and photographing for documenting hydrosalpinx, the same mouse genital tissues were fixed in 10% neutral formalin, embedded in paraffin and serially sectioned longitudinally (with 5 μm/each section). Efforts were made to include the cervix, both uterine horns and oviducts, as well as lumenal structures of each tissue in the same section. The sections were stained with hematoxylin and eosin (H&E). Three representative sections separated by 15μM or more from each other were used for evaluating oviduct dilation. The oviduct dilation was assessed under a 4x subjective lens. The severity of oviduct dilation was semi-quantitatively scored using the following criteria: 0, no significant oviduct lumenal dilatation; 1, mild dilation of a single cross section; 2, one to three dilated cross sections; 3, more than three dilated cross sections; and 4, confluent pronounced dilation. The median of the three scores served as a single value for each oviduct unilateral dilation score. Both unilateral scores for each mouse were combined to form a bilateral dilation score. Oviduct dilation incidence was calculated as the number of mice with a score of 1 or higher divided by the total number of mice in the group.

### 7. Statistics analyses

The time courses of live organism shedding (IFUs) were compared using “area-under-the-curve or AUC” between two groups using Wilcoxon rank sum test. The individual data point IFUs and pathology scores were also analyzed with Wilcoxon rank sum test. The category data including the % of mice positive for live organism shedding or pathology were analyzed using Fisher’s Exact test. Prior to any two-group comparison, an ANOVA was used to determine whether there was an overall significant difference among all groups in the same experiment (https://goodcalculators.com/one-way-anova-calculator/).

## Acknowledgement

This study is supported in part by US NIH grants (R01AI047997 & R56AI168479 to GZ)

